# A mixed generative model of auditory word repetition

**DOI:** 10.1101/2022.01.20.477138

**Authors:** Noor Sajid, Emma Holmes, Lancelot Da Costa, Cathy Price, Karl Friston

## Abstract

In this paper, we introduce a word repetition generative model (WORM), which—when combined with an appropriate belief updating scheme—is capable of inferring the word that should be spoken when presented with an auditory cue. Our generative model takes a deep temporal form, combining both discrete and continuous states. This allows a (synthetic) WORM agent to perform categorical inference on continuous acoustic signals, and—based on the same model—to repeat heard words at the appropriate time. From the perspective of word production, the model simulates how high-level beliefs about discrete lexical, prosodic and context attributes give rise to continuous acoustic signals at the sensory level. From the perspective of word recognition, it simulates how continuous acoustic signals are recognised as words and, how (and when) they should be repeated. We establish the face validity of our generative model by simulating a word repetition paradigm in which a synthetic agent or a human subject hears a target word and subsequently reproduces that word. The repeated word should be the target word but differs acoustically. The results of these simulations reveal how the generative model correctly infers what must be repeated, to the extent it can successfully interact with a human subject. This provides a formal process theory of auditory perception and production that can be deployed in health and disease. We conclude with a discussion of how the generative model could be scaled-up to include a larger phonetic and phonotactic repertoire, complex higher-level attributes (e.g., semantic, concepts, etc.), and produce more elaborate exchanges.

## Introduction

Word repetition—a deceptively simple language task—requires a person to reproduce a previously heard word spoken by someone else (Hanley et al., 2002; Hanley et al., 2004). It involves both the perception and production of the heard word, along with a contextual understanding of the linguistic exchange. Functionally, it can be viewed as deep inference, organised over nested time scales that successively unfold to generate auditory streams from ordered sequences or recognise auditory objects in continuous streams. The requisite deep or hierarchical organisation can be separated into two levels that interact with each other to repeat heard words. At the higher, slower level, word repetition requires turn-taking (Friston and Frith, 2015; Levinson and Torreira, 2015; Friston et al., 2020a): inferring when to listen for the auditory signal and when to respond. Whilst at the lower, faster level (i.e., within each turn), repetition requires some representation of the acoustic stream. When it is someone else’s turn to speak, the faster level assimilates incoming continuous acoustic signals; when speaking, it requires the generation of articulatory activity that recapitulates the previously heard word. In this way, turn-taking can be considered as inferring the context about whether one should be speaking or listening.

In this paper, we introduce a **wo**rd **r**epetition **m**odel (WORM) that instantiates this deep temporal organisation. To do this, we cast word repetition as a process of active inference (Friston et al., 2017c; Da Costa et al., 2020a; Friston et al., 2020b), which treats belief updating as a gradient descent on variational free energy (Hinton and Zemel, 1993), analogous to the evidence lower bound (Winn and Bishop, 2005). This is equivalent to maximizing the sensory evidence (a.k.a., marginal likelihood) for the generative model of how external (hidden) states or causes generate the sampled auditory signal (Friston et al., 2010). Technically, this implies maximising the marginal likelihood for our model of the sensed world or, more succinctly, self-evidencing (Hohwy, 2016).

In terms of the deep temporal organisation, WORM is furnished with discrete and continuous states that interact with each other to repeat the heard word at the appropriate time. The discrete states equip the model with the capacity to perform categorical inference over the incoming auditory signal, i.e., the recognition of the heard words. Equally, the continuous states allow the model to recognise (and produce) continuous auditory signals attached to some discrete label. This type of hierarchical interaction speaks to a mixed model with a slow-evolving discrete level, and a fast-evolving continuous level that facilitates inferences about the causes of particular (sampled) auditory observations. Happily, previous active inference mixed models have detailed the exact machinery required for this kind of integration of discrete and continuous levels (Friston et al., 2017c). WORM employs this particular active inference formulation, where higher-level discrete levels induce a short continuous auditory trajectory at the level below. This allows WORM to perform categorical inference on (longer) continuous acoustic signals, and to repeat heard words based on categorical states.

Our work builds on a long line of existing work within the domain of speech recognition and production. Previously, the focus has been on treating speech recognition and production as a learning problem based on either deep learning (Chorowski et al., 2014; Chan et al., 2016; Battenberg et al., 2017; Kim et al., 2017; Prabhavalkar et al., 2017; Chiu et al., 2018) or hybrid methods combining hidden Markov models (HMM) and neural networks (Bourlard and Morgan, 1994; Young et al., 1994; Senior et al., 2014). These approaches to speech recognition limit themselves to learning associations between the input and output via the training of a neural network architecture. We provide a brief overview of current approaches for computational modelling of word repetition in Table 1; these deal with both recognition and production of the auditory signal.

**Table 1.**
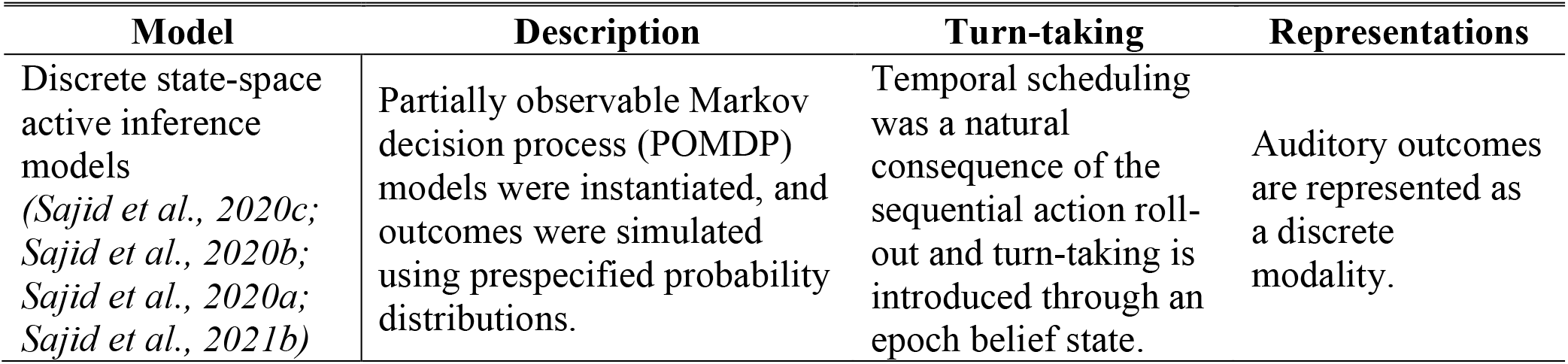
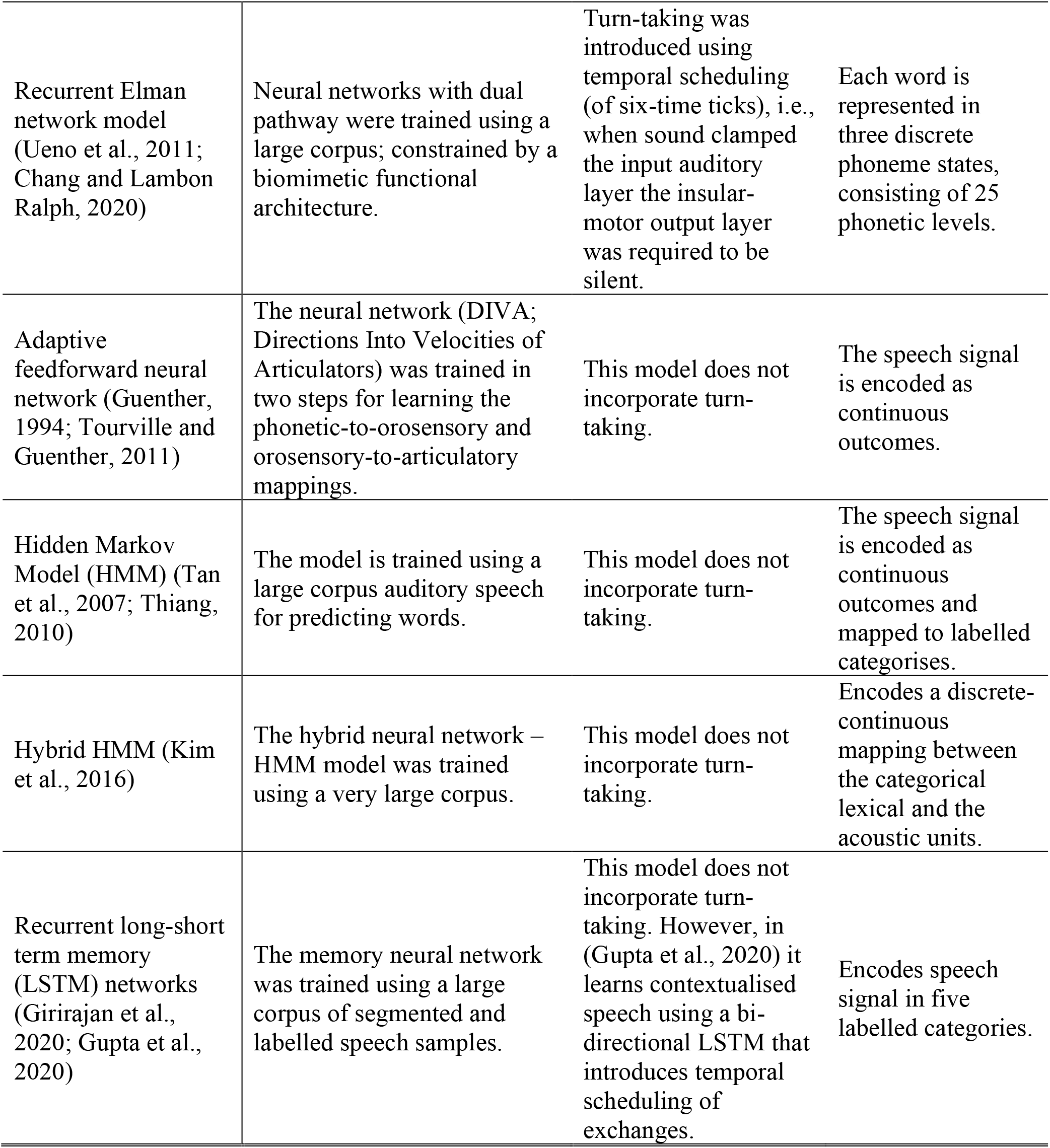
Recent computational models of word repetition. Each row summarises a different type of model. The columns briefly describe the models, how they deal with turn-taking, and the types of representations they use (e.g., discrete or continuous).

| Model | Description | Turn-taking | Representations |
| --- | --- | --- | --- |
| Discrete state-space active inference models<br>(*Sajid et al., 2020c*;<br>*Sajid et al., 2020b*;<br>*Sajid et al., 2020a*;<br>*Sajid et al., 2021b*) | Partially observable Markov decision process (POMDP) models were instantiated, and outcomes were simulated using prespecified probability distributions. | Temporal scheduling was a natural consequence of the sequential action roll-out and turn-taking is introduced through an epoch belief state. | Auditory outcomes are represented as a discrete modality. |
| Recurrent Elman network model (Ueno et al., 2011; Chang and Lambon Ralph, 2020) | Neural networks with dual pathway were trained using a large corpus; constrained by a biomimetic functional architecture. | Turn-taking was introduced using temporal scheduling (of six-time ticks), i.e., when sound clamped the input auditory layer the insular-motor output layer was required to be silent. | Each word is represented in three discrete phoneme states, consisting of 25 phonetic levels. |
| Adaptive feedforward neural network (Guenther, 1994; Tourville and Guenther, 2011) | The neural network (DIVA; Directions Into Velocities of Articulators) was trained in two steps for learning the phonetic-to-orosensory and orosensory-to-articulatory mappings. | This model does not incorporate turn-taking. | The speech signal is encoded as continuous outcomes. |
| Hidden Markov Model (HMM) (Tan et al., 2007; Thiang, 2010) | The model is trained using a large corpus auditory speech for predicting words. | This model does not incorporate turn-taking. | The speech signal is encoded as continuous outcomes and mapped to labelled categorises. |
| Hybrid HMM (Kim et al., 2016) | The hybrid neural network – HMM model was trained using a very large corpus. | This model does not incorporate turn-taking. | Encodes a discrete-continuous mapping between the categorical lexical and the acoustic units. |
| Recurrent long-short term memory (LSTM) networks (Girirajan et al., 2020; Gupta et al., 2020) | The memory neural network was trained using a large corpus of segmented and labelled speech samples. | This model does not incorporate turn-taking. However, in (Gupta et al., 2020) it learns contextualised speech using a bi-directional LSTM that introduces temporal scheduling of exchanges. | Encodes speech signal in five labelled categories. |

Conversely, WORM seeks to understand whether—by framing speech recognition as an active (Bayesian) inference problem with an underlying generative model—we can infer the causal relationships between input and output, thereby gaining a structural understanding of the sequences of words being presented and their context sensitivity. Unlike previous models (see Table 1), WORM is a mixed generative model that incorporates both continuous and discrete states. The generative model has higher-level discrete layers that map to a lower-level continuous auditory time series. Briefly, the lower level contends with speech recognition and production, i.e., categorical inference from acoustic signals and production of acoustic signals from categorical states. Accordingly, it can receive both a continuous acoustic signal and, phonetically understand the heard speech, by parsing the continuous signal into discrete segments (i.e., speech segmentation). The higher level contends with turn-taking, based on a narrative about whose turn it is to speak at a given time (Mirza et al., 2016; Friston et al., 2020a).

This equips WORM with the capacity to simulate simple exchanges (observed in natural linguistic communication) in which the listener might be required to segment longer acoustic signals beyond simply hearing a single word. Accordingly, our model formulation extends the standard word repetition experimental scenario, e.g., (Swinburn et al., 2004); where the participant is instructed what to do before the task begins, hear a single word and repeat it back. This extension is relevant for clinical and experimental settings in which the experimenter-subject interactions might involve natural exchanges that deviate from the standard paradigm; for example, as the subject requests a repetition of the target word.

The remaining paper is structured as follows. We first briefly introduce active inference and mixed generative models. Next, we set out the problem of word repetition and introduce our new generative model (i.e., WORM), which specifies how (and when) a continuous acoustic signal is generated, given a sequence of words with discrete attributes. The third section establishes the face validity of WORM, by simulating word repetition. We show that a WORM agent is capable of correctly repeating a heard word, in a manner that is not acoustically identical to the heard word—and can interact naturally with either another WORM agent or with a human subject, taking its turn to listen and speak at appropriate times. We conclude with a brief discussion of applications and future extensions of WORM.

## Active inference and mixed generative models

Briefly, active inference characterises the brain as an inferential, self-evidencing system that infers the causes of sensory samples while, at the same time, acting to solicit sensations that are the least surprising. Formally, this requires the minimisation of current and future surprisal (i.e., variational and expected free energy) about current and future observations (Friston et al., 2017b; Da Costa et al., 2020a; Sajid et al., 2021a), given a probabilistic generative model describing how external states cause sensations. Under this framework, one can generate distinct behaviours using different generative models. For example, active inference has been shown to successfully simulate a wide range of complex behaviours, including word repetition with fully discrete models (Sajid et al., 2020a; Sajid et al., 2020b; Sajid et al., 2021b), dyadic exchanges (Friston and Frith, 2015; Friston et al., 2020a), active listening (Friston et al., 2020b), active vision (Parr et al., 2021) and scene construction (Friston et al., 2017d; Parr and Friston, 2017; Heins et al., 2020).

What follows is a detailed description of the generative model used to illustrate the belief updating and the subsequent performance of synthetic subjects. Generally speaking, getting the generative model right is the most important step in constructing a plausible account of sentient behaviour. Given a generative model, realistic behaviour can be reproduced by inverting the model using standard (variational) message passing schemes (Friston et al., 2017a). Crucially these schemes have a degree of biological plausibility, to that the extent they can reproduce neuronal dynamics, their electrophysiological correlates and accompanying behaviour (here, choosing what to say and then articulating it in real-time).

In specifying the generative model, one has to consider all of the latent states and contingencies that are necessary to generate sensory cues (and behaviour) that characterise a particular task or paradigm. This specification can be regarded as instantiating the requisite intentional or cognitive set— i.e., explaining to the subject what is expected of them—by installing priors and causal architectures that are apt for the paradigm at hand.

Compared to previous (discrete state-space) active inference word repetition models, mixed generative models have several advantages including i) recognising and simulating continuous auditory signals from discrete categories, ii) being able to verbally communicate with the model in real-time and iii) computational phenotyping of recorded subject behaviour (Schwartenbeck and Friston, 2016).

These mixed generative models integrate distinct, hierarchically composed, levels of discrete and continuous states. The (higher) discrete level pertain to discrete outcomes (e.g., a heard word) caused by discrete hidden states (e.g., a belief about the identity of the target word) (Friston et al., 2017b). Figure 1 illustrates a general form of such discrete models; specifically, a partially observable Markov decision process. Briefly, outcomes (*o*) and hidden states (*s*) are generated by three sets of categorical probability distributions, parametrised by *A, B* and *D*. The first distribution, *A*, is the likelihood, which maps hidden states to outcomes. The second, *B*, is associated with the probabilistic transitions among hidden states. Lastly, *D* specifies the probabilities of initial states. Policies, *π* (i.e., sequences of actions or control states *u*) specify transitions among some states, which in turn generate expected outcomes (e.g., a heard word). Under active inference, actions are selected to minimise the surprisal of expected outcomes (*G*) ; e.g., saying or hearing the word ‘sun’ after ‘bright’, as opposed to ‘chocolate’.

**Figure 1.**
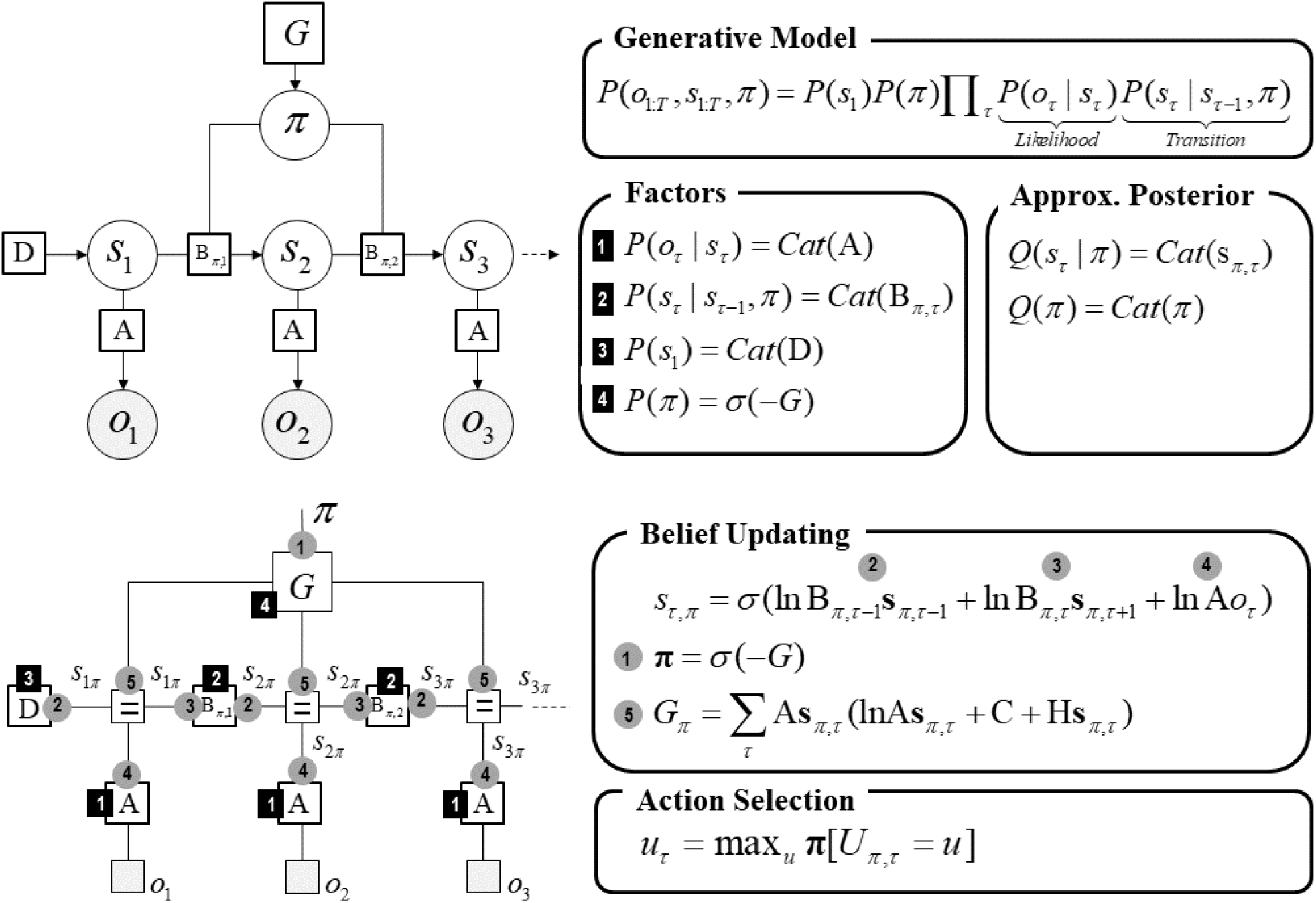
*A generic generative model for discrete states and outcomes: adapted from* (Friston et al., 2017c). The upper panel presents a Bayesian network representation of the model, which depicts the conditional dependencies among hidden states and how they cause outcomes. Here, open circles are random variables, filled circles denote outcomes and squares indicate known variables. The accompanying equations specify the generative model—a joint probability of outcomes and their causes. The model is expressed in terms of a likelihood, transition function, and priors over causes. The likelihood is specified by a matrix A whose elements are the probabilities of an outcome under every combination of hidden states. The probabilistic transitions specified in matrix B depend upon actions, which are determined by policies (π). *Cat* denotes a categorical probability distribution. The key aspect of this generative model is that policies are more probable a priori if they minimise the (time integral of) expected free energy G (i.e., expected surprisal), which depends upon prior preferences (encoded by C) and the uncertainty about outcomes under each state (encoded by H). Finally, the vector *D* specifies probabilities for each initial state. To estimate the hidden states—and other variables that cause outcomes—we need to invert the model through variational Bayesian inference. Variational inference requires specification of a family of approximate posterior distributions. For this, we employ a mean-field approximation, in which posterior beliefs are approximated by the product of marginal distributions across time points. The lower left panel is the equivalent representation of the Bayesian network (upper left panel) in terms of a Forney factor graph. Here, the nodes correspond to factors and the edges to unknown variables. Filled squares denote observable outcomes. The edges are labelled in terms of the sufficient statistics of their marginal posteriors. Factors have been labelled in terms of the parameters encoding the associated probability distributions and the circled numbers correspond to the messages that are passed from nodes to edges. These correspond to the messages implicit in the belief updates. The accompanying equalities are the belief updates mediating approximate Bayesian inference and action selection; for technical details see (Friston et al., 2017c; Da Costa et al., 2020a).

Similarly, mixed models generate continuous signals from discrete representations of external states. For our purpose, we use the mixed generative model of spoken word sequences introduced in (Friston et al., 2020b) (Figure 2). This model includes various representations that effect the generation an acoustic signal, which are labelled as lexical, speaker, and prosody states. The model generates acoustic signals from these discrete states. Lexical states control aspects of the acoustic signal related to which word is spoken, speaker states control aspects related to a person’s voice, and prosody states control aspects related to how a word is spoken (e.g., a happy or sad tonal inflexion). Briefly, the generative model constructs a time-frequency representation based on the lexical content of the word, which is transformed into distinct transients that incorporate within-transient prosodic inflexions. Finally, these transients are aggregated into a continuous time series, corresponding to the acoustic signal. Word recognition then corresponds to the inversion of this generative model: for the ensuing categorical inference based on acoustic signals, we employ *active listening* (Friston et al., 2020b). This entails covert action selection that determines the placement of word boundaries. The ‘active’ component is the placement of word boundaries at particular positions within the continuous acoustic signal. For each possible word interval, the likelihoods of model parameters are evaluated—and the interval with the greatest evidence is selected. For technical details, see (Friston et al., 2020b).

**Figure 2.**
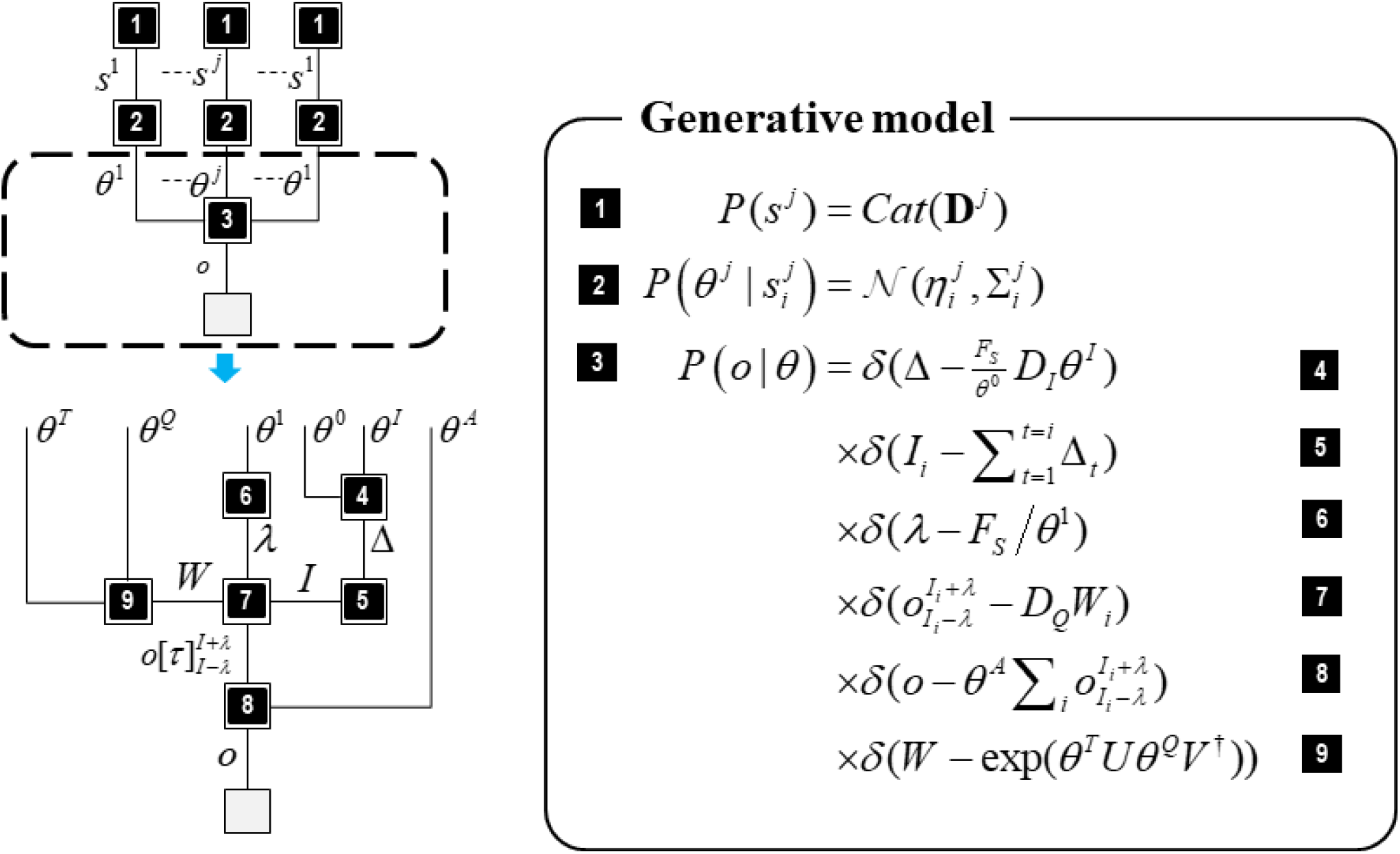
*A mixed generative model for synthesising and recognising speech, adapted from* (Friston et al., 2020b). The left panel represents a Forney factor graph of the continuous model, denoted by the factorised probability densities that underwrite the generative model. The factors are indicated by numbered squares, and the edges (i.e., lines connecting the factors) represent the variables common to the connected factors. Here, *s* are the discrete hidden states and 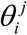 are sampled parameters from state-dependent distributions, *o* the continuous outcome, *θ* ^*Q*^ discrete cosine transforms, *W* is the time-frequency representation of the spoken word, Δ*I* transients, *I* the fundamental interval variable, *θ* ^0^ the average fundamental frequency, *θ* ^*I*^ the inflexion around the average fundamental frequency, *θ* ^*T*^ the inverse temperature parameter, *θ* ^*A*^ amplitude parameter. The top graphic illustrates factors 1–3, and the bottom graphic unpacks factor 3 in terms of factors 4–9. The process of generating data requires a series of local operations taking place at each factor from top to bottom i.e., sample states from factor 1, then parameters from factor 2, then perform the series of operations in factor 3 to construct the time-series. The inversion of this model entails bidirectional message passing across each factor node, such that empirical priors and likelihoods are combined at each edge to form posterior beliefs about the associated variable. The second panel formalises each factor. Factor 1 is the prior probability associated with the hidden states and takes a categorical form. Factor 2 is a normal distribution that specifies the dependence of parameters on states. Note, that each discrete state is associated with a distinct expectation and covariance for the parameters. Factor 3 describes how the observed time-series is generated from the parameters, and this is decomposed into factors 4–9. Factor 5 determines the internal ‘action’ that selects the interval for segmentation.

Our particular mixed model generates a continuous acoustic signal corresponding to short spoken phrases by integrating the two distinct model formulations: (i) mappings between discrete states and outcomes (Figure 1) and (ii) (amortised) mappings from discrete states to continuous outcomes (Figure 2). This entails a deep temporal organisation, where the slowly evolving discrete states contextualise the linguistic exchange, and fast continuous states facilitate inferences about the causes of (sampled) auditory observations (Friston et al., 2017c). Formally, the discrete outcomes specify a fixed-point attractor for inferences over continuous acoustic signals (Friston et al., 2017b) and only the discrete model can determine state transitions that enable periodic switching of hidden causes generating dynamics. This is required for our (turn-taking) word repetition paradigm, where each turn requires the recognition (or production) of specific sequences of words that only update at the start of the next ‘turn’ or epoch of exchange.

Belief updating in these types of mixed models operates as follows: (i) descending messages comprise Bayesian model averages of predicted outcomes and (ii) the ascending messages from the lower, continuous, level of the model are the posterior estimate of these outcomes, having sampled some continuous observations (Friston et al., 2017c). Explicitly, descending messages dictate the empirical priors over the dynamics of the lowest (continuous) level that returns the corresponding posterior distribution over priors given the continuous data at hand. For technical details about the inferential scheme used to invert mixed generative models see (Friston et al., 2017c; Friston et al., 2020b).

Practically, we introduced an amortised form of the continuous active listening model introduced in (Friston et al., 2020b). Specifically, we used the active listening scheme during the pre-training phase (see below) to estimate the model parameters over the lexical, speaker, and prosody states for a particular set of words. These parameter estimates were used as empirical priors in our generative model. Importantly, this allowed us to focus on making inferences about, and produce, short spoken phrases (i.e., sequences of words), and ensured computational simplicity to simulate real-time interactions between two agents. This meant that word segmentation was learned, as opposed to being inferred online, under the Bayesian beliefs afforded by the discrete part of the model. In effect, we assumed that the interlocutors were sufficiently familiar with each other that they could recognise individual words, without having to resolve any ambiguity about word boundaries. Practically, this meant that the agent was trained using slow and clear speech, with unambiguous word boundaries.

## WORM: a word repetition model

Word repetition involves both the perception and production of an acoustic signal, along with a contextual understanding of turn-taking in the exchange. In natural speech, no two instantiations of a particular spoken word are acoustically identical. Yet, categorical perception means that we can extract a word category from an acoustic signal, which ultimately allows us to understand the semantic content of speech. When humans perform a word repetition task, the acoustic response is not identical to the target word— yet it is considered to be correct if the lexical category is the same.

To simulate word repetition, a generative model requires both discrete and continuous states: continuous states that represent the acoustic signal and discrete states that represent the word category. The mapping from discrete word states to continuous acoustic states has two purposes: From the perspective of speech perception (i.e., Bayesian model inversion), it enables categorical inference of the incoming auditory signal, i.e., to identify the heard word. From the perspective of speech production, it enables the model to generate continuous auditory signals, corresponding to a discrete word label, regardless of the precise acoustics of the heard target word (such as attributes related to the person’s voice). In addition, the word must be repeated at the appropriate time, which requires an inference about whether the person should be speaking or listening. This calls for a deep temporal model, in which words or phrases are present (either heard or repeated) over several timescales. In other words, the same phrase persists for the duration of a word sequence. And the same word persists for the duration of a phonemic sequence.

Rather than using individual words, most natural interactions involve sequences of words (i.e., phrases or sentences). For example, if we wanted someone to repeat a word, we would not simply utter the word and expect them to repeat it, but instead we might ask a question, such as “Can you please repeat triangle?” Similarly, in response, we might say something like “Ok, triangle” rather than simply stating the word alone. To simulate these types of interactions, we use phrases—that comprise sequences of words—in our model. This requires us to specify how words transition over time (which can be thought of as syntax).

WORM is illustrated in Figure 3. It contains two discrete levels. The higher level maps discrete states to discrete outcomes. This level has three (hidden) state factors (*Context, Heard word*, and *Spoken word*) and three outcome factors (*Target Word, Feedback*, and *Syntax)*. The *Context* factor contextualises the linguistic narrative: being asked to repeat the target word (i.e., Question), responding to the question and repeating the heard word (i.e., Answer), and receiving performance evaluation (i.e., Feedback). The *Heard Word* factor has states corresponding to a heard word: Green, Red, Blue, Triangle, or Square. The *Spoken Word* states cover what could be spoken: Green, Red, Blue, Triangle, or Square. In terms of outcome modalities for this level, the *Target Word* outcome reports the heard or spoken word: Green, Red, Blue, Triangle, or Square. The *Feedback* outcome represents the positive or negative response received (only provided when *Context* is Feedback). The *Syntax* represents the sequences of words, with grammar used as absorbing states to indicate that the phrase has concluded. Here, syntactic structures are limited to one sort of question (e.g., ‘Ok, can you please repeat Green?’), 5 sorts of answers (e.g., ‘I am not sure’ or ‘Yes, Green’) and two sorts of feedback (“Correct!” or “Wrong!”).

**Figure 3.**
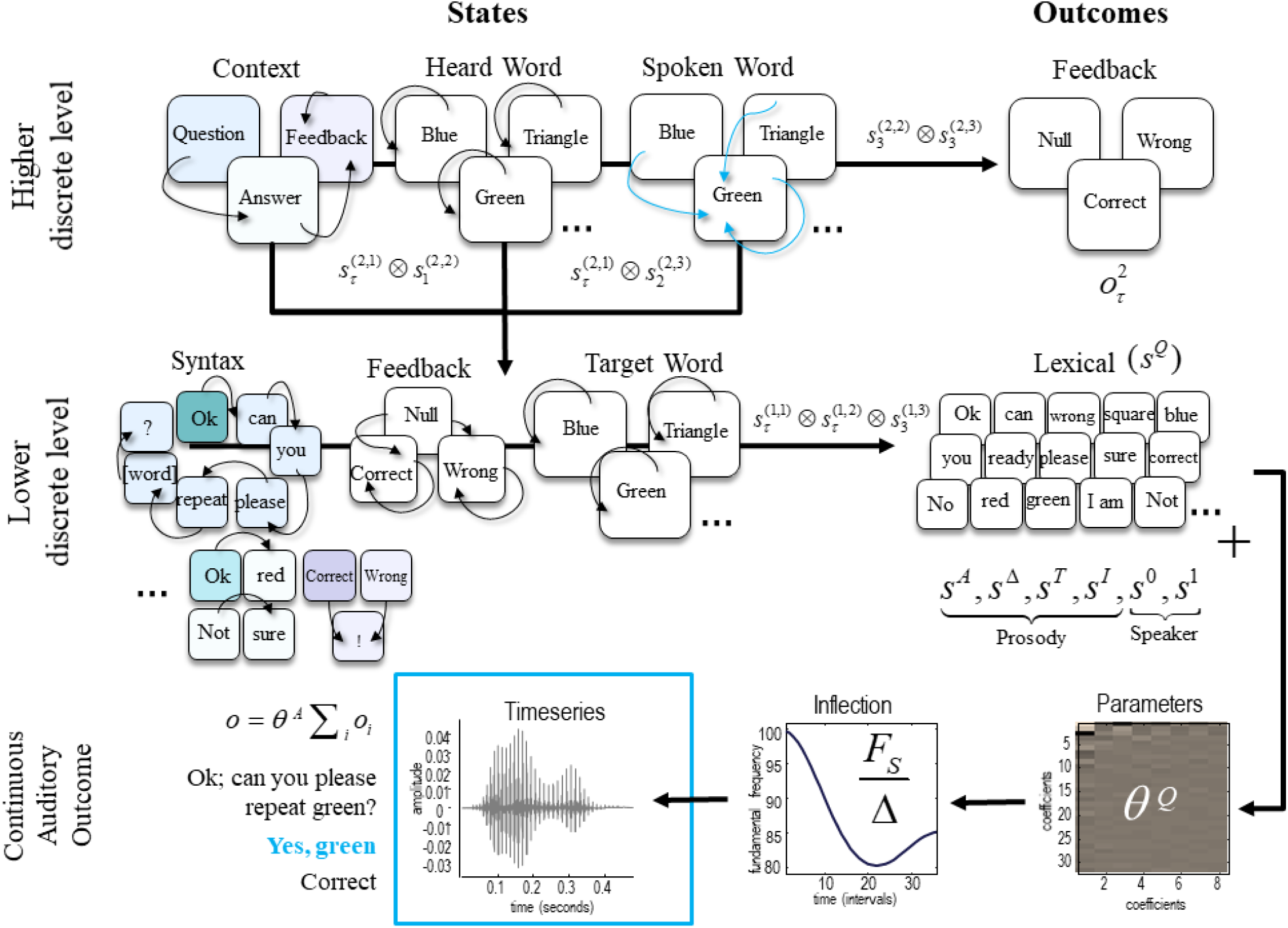
Word repetition model – WORM. This figure provides a schematic illustration of the generative model. The schematic displays the architecture involved in generating a linguistic exchange during the repetition of particular words, i.e., being asked to repeat a particular word, repeating it, and receiving feedback on the repetition. Briefly, this is a mixed generative model formulated in terms of discrete hidden states that map to a continuous auditory outcome. It has two levels of hidden states, where the higher (deeper) level unfolds slowly in time—furnishing contextual (turn-taking) constraints on the lower level that generates the continuous sequence of words. At the higher discrete level, transitions among *Context* states generate a sequence of phrases that have temporal order from Question, Answer and Feedback. The content of these phrases depends upon interactions with other hidden states in the model: if it is a Question, the *Spoken Word* is used in the question; if it is an Answer, the *Heard Word* is used in the answer. Each of these factors have five levels (which are not all displayed in the figure): blue, red, green, square or triangle. In this generative model, the *Spoken Word* state is policy-dependent; see the blue arrows that illustrate articulating the word ‘green’. In other words, policies determine transitions among controllable states, so that the *Spoken Word* states are selected intentionally and influence the *Target Word* at the level below. For example, when the Question contains the *Heard Word* Green, the *Spoken Word*—which is used in the Answer—is selected as Green. The combination of higher-level discrete states generates the *Syntax, Target Word*, and *Feedback* content (i.e., the states at the lower level). The hidden Syntax states at the lower level comprise sequences of words and grammar (i.e., ‘?’ and ‘!’) that make the exchanges more natural; for example, an Answer containing Green can be structured as “Ok, green”, “Sure, green” or “Yes, green”. The words encoded by the *Target Word* state are determined by the *Heard Word* factor when *Context* is Question and *Spoken Word* factor when *Context* is Answer. The first word of the phrase corresponds to the initial Syntactic state at the lower level, which is determined by the interactions among states at the higher level (encoded by darker shaded squares). For example, if the *Context* state is a Question, then the initial syntax state is the word ‘Ok’, no matter which of the five *Heard Word* states are selected at the higher level. The transition matrices at the lower discrete level then determine subsequent words (black arrows), by specifying transitions among *Syntax* states that depend upon the *Context* states at the higher level. If the *Context* state is Answer, then the initial *Syntax* state is ‘Yes’, ‘No’, ‘I am’, ‘Sure’, ‘Ok’ depending upon high order interactions among the remaining high-level states: a “Yes” is generated when *Heard* and *Spoken word* match. For brevity, Syntaxes that generate antonymous phrases—although defined within the model—have been omitted from the graphic; for example, a ‘Not sure’, ‘No’, ‘I am not ready’ answer. The next stage is to map discrete states at the lower-level to continuous outcomes at each time step of the process. These outcomes are single words. For this, we employ an active listening model for synthesising and recognising speech (Friston et al., 2020b). In brief, this involves using these empirical priors to (i) construct a time-frequency representation based on the lexical content and random assignment of speaker and prosody parameters, (ii) transform them into distinct transients that incorporate within-transient prosodic inflexions, and (iii) aggregate and scale the transients to construct the auditory time series.

In Figure 3, the arrows that link states within each factor model their transitions over time (B). These encode prior beliefs about trajectories. Transitions within the *Context* factor dictate the type of narrative currently in play. In this model, Question transitions to Answer and Answer transitions to Feedback. A linguistic exchange terminates when feedback has been received, i.e., feedback is an absorbing state. For the *Spoken word* factor, there are five possible transitions, each to one of the five possible words, dependent upon the selected action. For example, when the word “Green” is chosen for repetition, the state transitions to Green (highlighted in Fig. 3), irrespective of the previous word (red, triangle, etc.). In contrast, the *Heard word* transition matrix is an identity matrix which means that the *Heard word* stays the same during an exchange. The arrows between states and outcomes represent the likelihood distribution (A). The *Feedback* likelihood depends on all the hidden states. Correct feedback is given at the final exchange, if the previously heard word is repeated correctly, and Wrong feedback is given otherwise. When the *Context* is Question or Answer, Null feedback is given, irrespective of the *Heard* and *Spoken word*. The *Target Word* likelihood depends on either *Spoken word* (when the *Context* is Answer) or *Heard word* (when the *Context* is Question). The likelihood is defined as a one-to-one mapping between the *Heard word* and *Target Word* if the context is Question. Otherwise, there is a one-to-one mapping between *Spoken word* and *Target Word*. The *Syntax* likelihood depends on the *Context*. For example, Question *Context* constrains the syntax outcomes to ‘Ok’, ‘Can’, ‘You’, ‘Please’, ‘Repeat’.

The lower level encodes how the discrete outcomes from the level above evolve over time. The initial *Syntactic* state at this level is determined by the interactions among states at the higher level, encoded by the mapping *D*. For example, if the *Context* state is a ‘Question’, then the initial *Syntax* state is the word “Ok”, and the transitions then determine subsequent words (Figure 3; illustrated by the black arrows), by specifying transitions among *Syntax* states that depend upon the Question and *Heard word* states at the higher level. However, if the *Context* state is Answer, then the initial syntax state can be Yes, No, Ok, I am, or Sure, depending upon high order interactions among the high-level states. For the *Context* state ‘Feedback’, the initial Syntax state depends on the alignment of the *Heard and Spoken word* state: ‘Correct’ will be generated when these two states match and ‘Wrong’ otherwise. The initial *Syntax* states determine the plausible state transitions at this level. For example, if the *Context* state is a Question, then Ok will be followed by ‘can you please repeat [word]?’. Here, the particular word is determined by the *Heard Word*. These particular discrete states are mapped to continuous outcomes at each time step of the generative process using an amortised form of the speech production and recognition generative model introduced in (Friston et al., 2020b) (Figure 2) with pre-specified empirical priors.

## Simulations

Here, we present two sets of *in silico* ‘experiments’ that underwrite the face validity of WORM. These involve exchanges between a WORM model (“WORM 1”) and another WORM model (“WORM 2”; control) (Table 2; Simulation ID Z) and/or a human (Table 2 Simulation ID A-C). Experiment Z is the control experiment between two WORM agents. In contrast, experiments A-C evaluate the WORM’s capacity to interact with a human subject. Here, we deliberately introduced the human subject at different stages of the trial to ask the question, to answer and provide feedback. For all experiments, the lexical content, temporal duration (i.e., six epochs) and human subject were kept consistent. The human subject was female, 27 years old and had normal hearing.

**Table 2.** Simulation set-up

| Simulation ID | Type | Agents (Question – Answer – Feedback) |
| --- | --- | --- |
| Z | Control | WORM 1 – WORM 2 – WORM 1 |
| A | Human-in-the-loop | Human – WORM 1 – WORM 2 |
| B | Human-in-the-loop | WORM 1 – Human – WORM 1 |
| C | Human-in-the-loop | WORM 1 – WORM 2 – Human |

### Pre-training

For the simulations, we inverted a stream of continuous auditory signals articulating twenty-two distinct words into their discrete state-space representations. These discrete states were lexical, prosody and speaker. We used this to encode pre-specified empirical prior probability distributions, for each word, in our word repetition model. This can be regarded as introducing an amortised nonlinear mapping from discrete states to continuous auditory outcomes. To specify this, we followed the inversion process introduced in (Friston et al., 2020b). First, the auditory time series was segregated into transients. These were defined within the time intervals of the auditory stream and formant scaling and duration states defined. These time intervals were segmented using an active inference process. This determined the onsets and offsets of the interval containing a particular word from a set of possible boundary intervals and selected the interval with the greatest model evidence; see the description above or (Friston et al., 2020b) for technical details. The time-frequency representation was constructed from these transients, and the lexical parameters were evaluated from the constructed time-frequency representation.

These parameters were used to encode the *Lexical, Prosody* and *Speaker* attributes during the simulations. We estimated the parameters for the following words: ‘I am’, ‘OK’, ‘a’, ‘blue’, ‘can’, ‘correct’, ‘green’, ‘is’, ‘no’, ‘not’, ‘please’, ‘ready’, ‘red’, ‘repeat’, ‘sorry’, ‘square’, ‘sure’, ‘there’, ‘triangle’, ‘wrong’, ‘yes’, and ‘you’. The training data was collected from two different speakers and included each word being spoken 32 distinct times. The first speaker was female, 27 years old and had normal hearing, and second speaker was male, 62 years old and had normal hearing. This allowed us to capture the variability in natural speech, where each instantiation of a spoken word tends to be acoustically distinct.

### Diachronic inference and linguistic exchanges

To simulate linguistic or turn-taking exchanges, we rely upon the diachronic inference scheme introduced in (Friston et al., 2020a) (Figure 4). This scheme stipulates that each agent (i.e., simulated model) can either listen or speak at a given time point, to allow for a turn-taking exchange. This means that the beliefs of one agent are inferred by the other, via an exchange of outcomes. This aligns the linguistic exchange between the agents and ensures that posterior beliefs are consistent with what is heard, and, at the same time, the generated output is consistent with those beliefs. If two agents follow this process, their beliefs will align to ensure an appropriate linguistic exchange. For word repetition, this would entail one WORM agent (*i*) asking the other WORM agent (*j*) to repeat a particular word as WORM agent *j* listens. This is followed by WORM agent *j* responding with the answer as WORM agent *i* listens and evaluates whether the correct word has been spoken.

**Figure 4.**
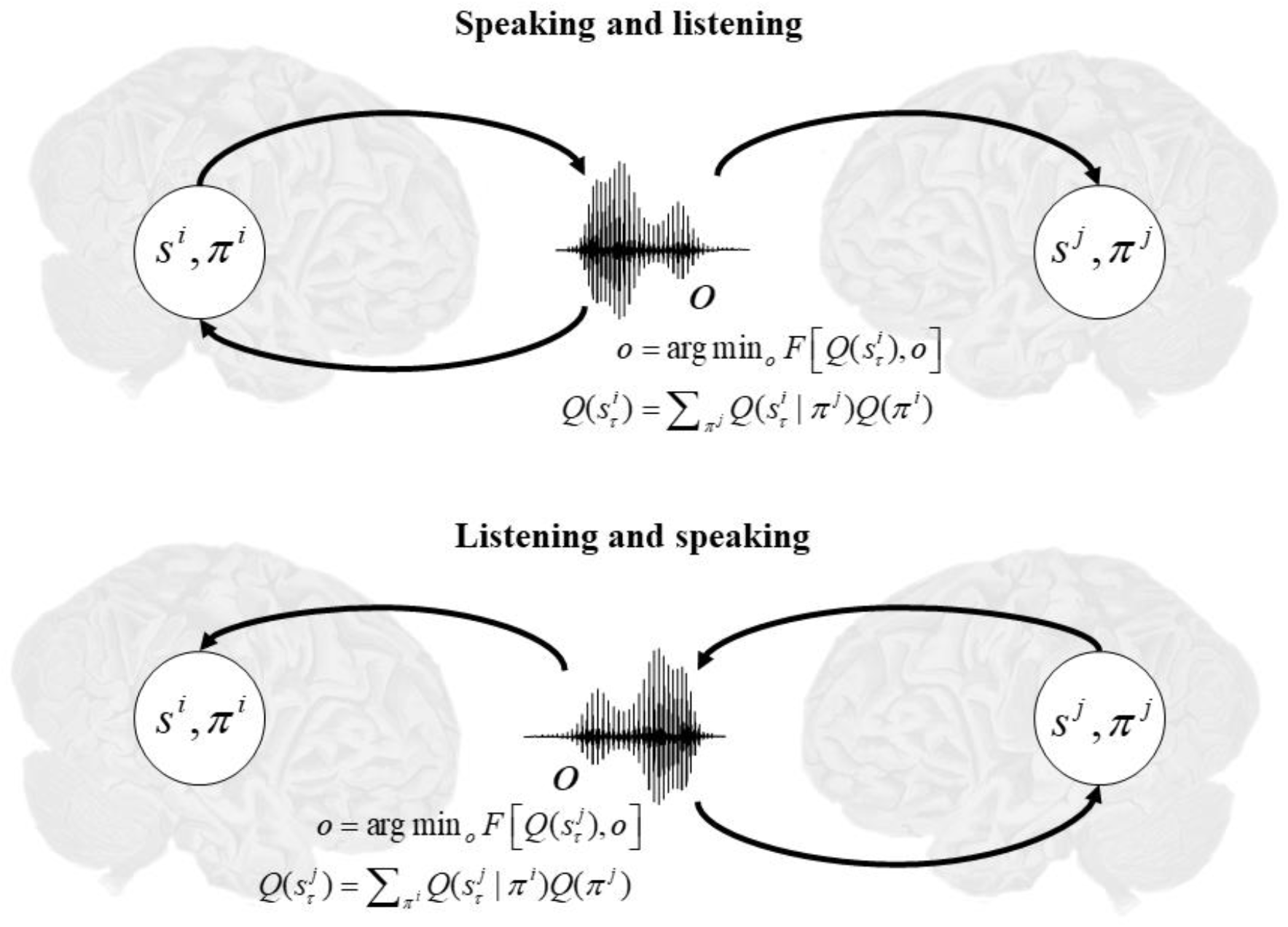
Pictorial representation of diachronic inference, *adapted from* (Friston et al., 2020a). In this setting, the outcomes (*o*) and actions (*u*) are assumed to have a one-to-one mapping. This is because actions generate outcomes that maximise model evidence. Furthermore, outcomes are shared between two models as they form the (Markov) boundary that separates the internal states of each model during the linguistic exchange. Accordingly, the internal states of one model (e.g., *i*) are the external states of another model (e.g., *j*) and *vice versa*. This scheme rests upon a turn-taking narrative, where only one model generates outcomes at any one time. Therefore, the models can only listen or speak at a given time point. The superscripts denote the two models (agents *i* and *j*). Briefly, these equations express how various states are sampled via the minimisation of the variational free energy. For technical details see (Friston et al., 2020a).

Next, we introduce how these particular agents can be another simulated WORM model or a real human subject interacting with WORM.

### Control simulations with two WORM models

First, we simulated 20 trials of a word repetition exchange between two WORM agents (Table 2: Simulation ID Z). Here, WORM 1 took on the role of an experimenter—asking for a particular word to be repeated and then evaluating how well that particular word was repeated. WORM 2 took on the role of the subject—appropriately repeating the target word. We repeated this simulation 20 times using different initialisation seeds for state estimation. This mimics the interaction of two different agents—with the same internal representations.

These simulations demonstrate that WORM can appropriately recognise and generate the target word during the linguistic exchange. During each trial, a different (random) target word was presented from the possible list of 5 words: Green, Red, Blue, Triangle, or Square. An example exchange, with the heard word “Square” is illustrated in Figure 5A. The accompanying belief updates for the *Heard Word* factor are shown in Figure 5B–D, each representing one time step at the higher level (corresponding to a Question, an Answer, then Feedback). Here, we can observe that inferring the heard word during the Question requires the entire sequence of words to be played out for the first 1.25 seconds (see Figure 5B). Conversely, during the Answer and Feedback, the model has appropriate expectations about the *Heard Word* from the beginning of the corresponding time step (see Figure 5C–D). This illustrates that the WORM agent can appropriately retain what was heard in previous time steps and generate outcomes consistent with those beliefs. During model evaluation, we observed 85% performance accuracy across 20 trials – validating the construction of the generative model. Importantly, the same WORM model can successfully encode target words based on different auditory streams. An example for ‘Square’ is presented in Figure 7.

**Figure 5:**
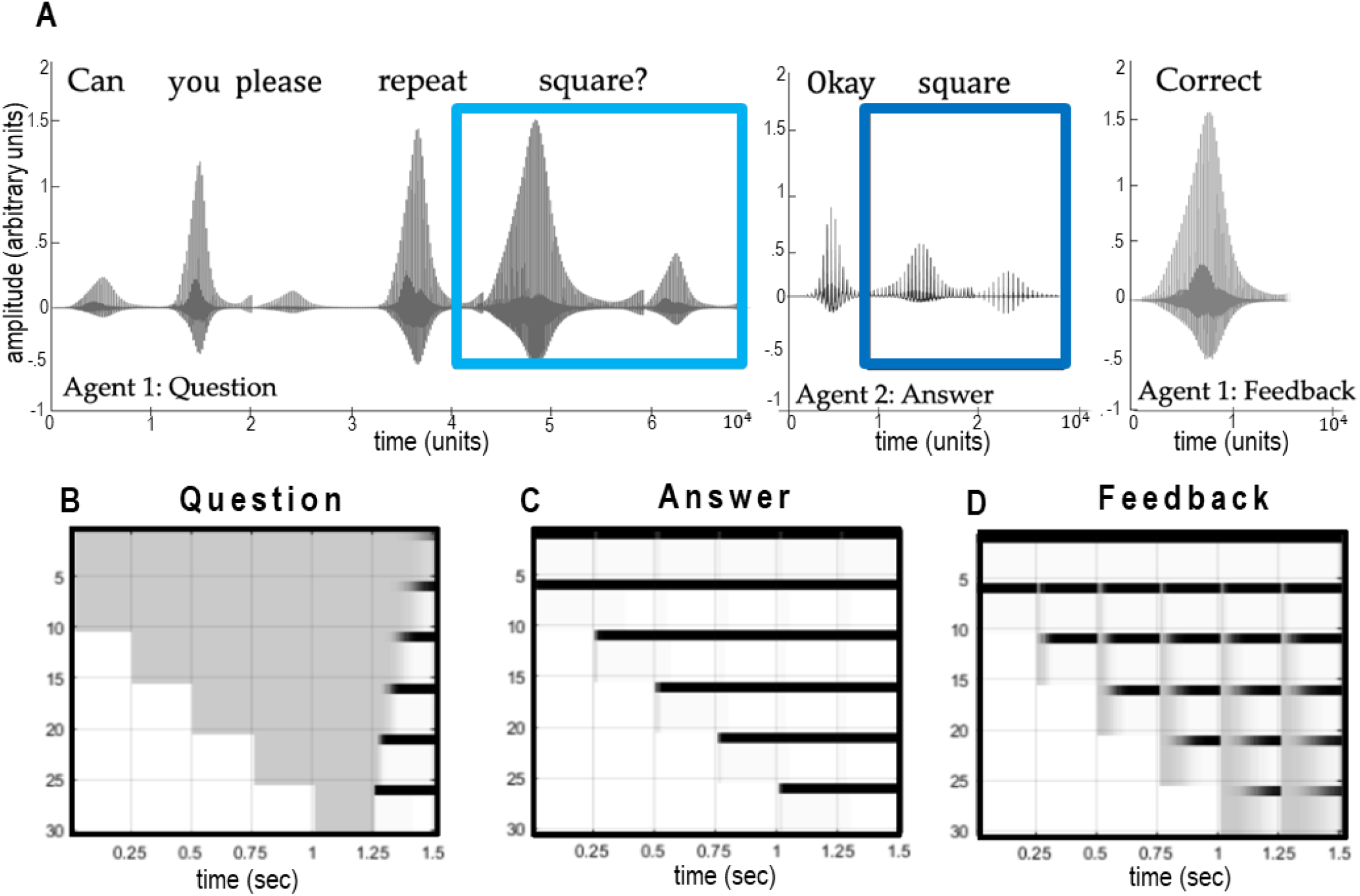
Control simulation. These graphics illustrate the simulation results from an example trial. Panel A presents the auditory waveforms produced during the word repetition exchange. The plots display the sound amplitude (y-axis) in the time domain (x-axis). Here, the question is asked by WORM 1, and the word “square” is indicated with a cyan rectangle. This is followed by WORM 2 responding with the word “square” (blue rectangle). Lastly, WORM 1 gives the feedback “correct”. Implicit in this sequence is the turn-taking exchange (or diachronic inference) that each agent can either listen or speak at a given time point. Panels B–D show the accompanying belief updates for the *Heard word* factor. Each panel reports belief updating over 6 epochs within each time step. The *x*-axis represents time in seconds (divided into 6 epochs), and the y-axis represents each of the associated states at different epochs (in the past or future). The shading represents posterior expectations. White is an expected probability of zero; black of one; and grey indicates gradations between these. Here, there are 5 states, and a total of 5 × 6 (states by epochs) posterior expectations. The first five rows in panel B correspond to expectations about the *Heard Word* in the first epoch, because there are five possible words. The second five rows are the equivalent expectations for the second epoch. This means that, at the beginning of the trial, the second five rows express beliefs about the future: namely, the next epoch. However, later in time, these beliefs refer to the past, i.e., beliefs currently held about the first epoch. This aspect of inference is effectively an implementation of working memory that enables agents to remember what has been heard—and accumulate evidence for the target word that is subsequently articulated. Note that most beliefs persist through time (along the *x*-axis). For example, the target word reveals itself after the first 5 epochs in panel B and this prospective belief is propagated into the future, and across exchanges.

**Figure 7:**
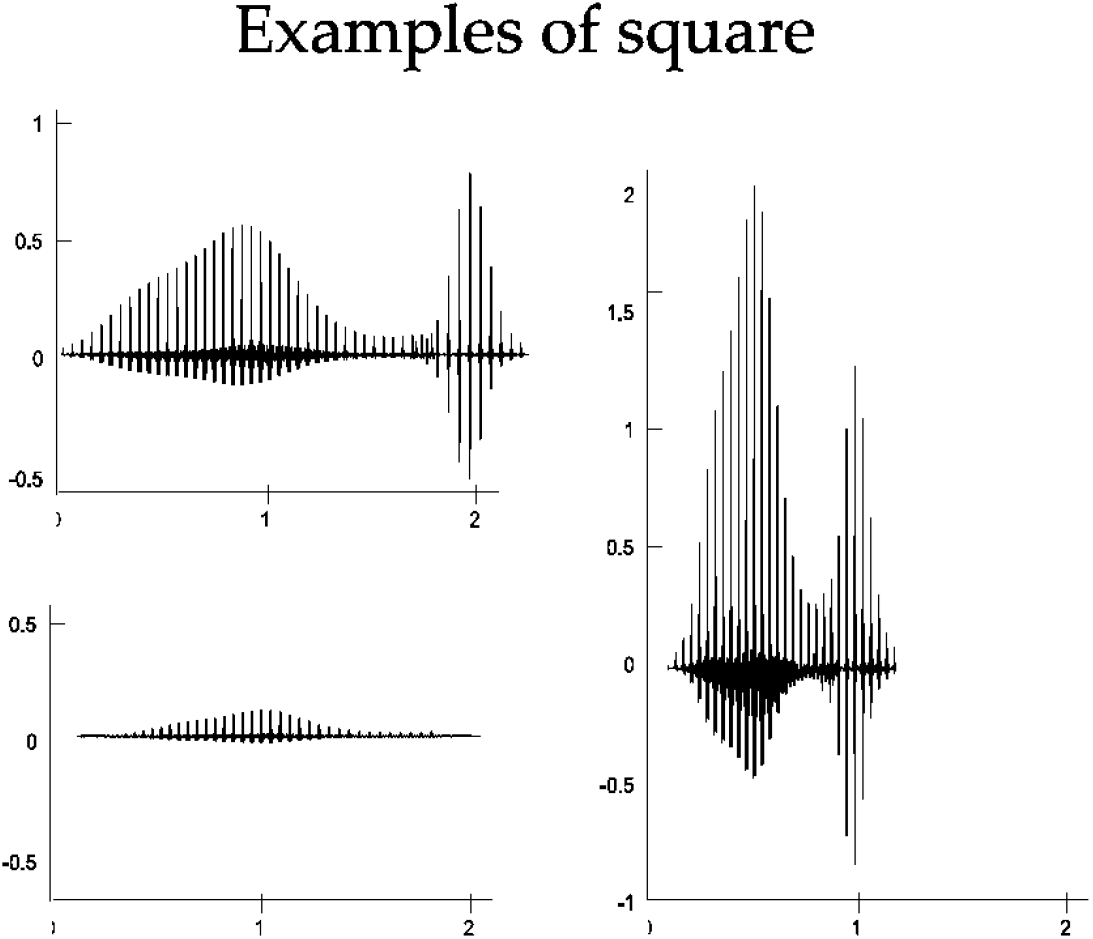
(Examples of auditory time series) This graphic plots a few examples of distinct waveforms generated by WORM for the word square. This simulates the natural variability of speech, where different instantiations of the same word tend to be acoustically distinct. Here, the x-axis denotes the time and the y-axis the sound amplitude.

### Simulations involving human interaction

Next, we assessed how WORM performs when interacting with a human subject. For this, we replaced one of the WORM models with a human when being asked to repeat a particular word. For two types of simulation, (Table 2: Simulation ID A & C), the human takes on the role of an experimenter and the WORM agent takes on the role of the subject. In this setup, the human asks for a particular word to be repeated, the agent repeats the target word and the human evaluates how well the word was repeated. Conversely, for the final type of simulation (Table 2: Simulation ID B), the WORM agent takes on the role of an experimenter, and the human takes the role of the subject having to repeat a heard word.

We simulated each experiment (A-C) for 20 different (random) target words from the possible list of 5 words: Green, Red, Blue, Triangle, or Square. The results show that WORM can appropriately recognise and generate the target word during the linguistic exchange with a human subject. We observed the following performance accuracy across the word repetition trials: For experiment A, during 80% of the trials the WORM agent’s response to the question matched the target word chosen by the human subject. For experiment B, during 80% of the trials the WORM agent was able to infer that the correct target word (and action) had been selected by the human subject. Lastly, for experiment C, during 95% of the trials the WORM agent was able to appropriately recognise the feedback articulated by the human subject (e.g., correct when correct was spoken). We observed that failure to correctly infer the continuous auditory signal was due to background noise (i.e., due to differences in microphone), similarity between particular words and inability to actively infer appropriate word boundaries.

We plot an example exchange when repeating the word ‘Blue’ in Figure 8; whether they are asking the question (Figure 8A), repeating the heard word (Figure 8B), or providing feedback (Figure 8C). It is important to note that WORM can effectively recognise the heard outcomes despite the noisy waveform produced by the real subject (Figure 8A), compared to the simulated articulation of the same phrase (Figure 8B). The ability to recognise words in continuous speech rests on using empirical priors—at the lower continuous level—acquired with the same human voice. As in the control simulations in Figure 5, WORM requires the entire sequence of words to be played out for the first 1.25 seconds to identify the target word during the Question (Figure 8A). It can also recognise that the action selected (i.e., the word being spoken by the real subject is ‘Blue’; Figure 8B: final panel), and the feedback required for a correct response (Figure 8C: final panel).

**Figure 8.**
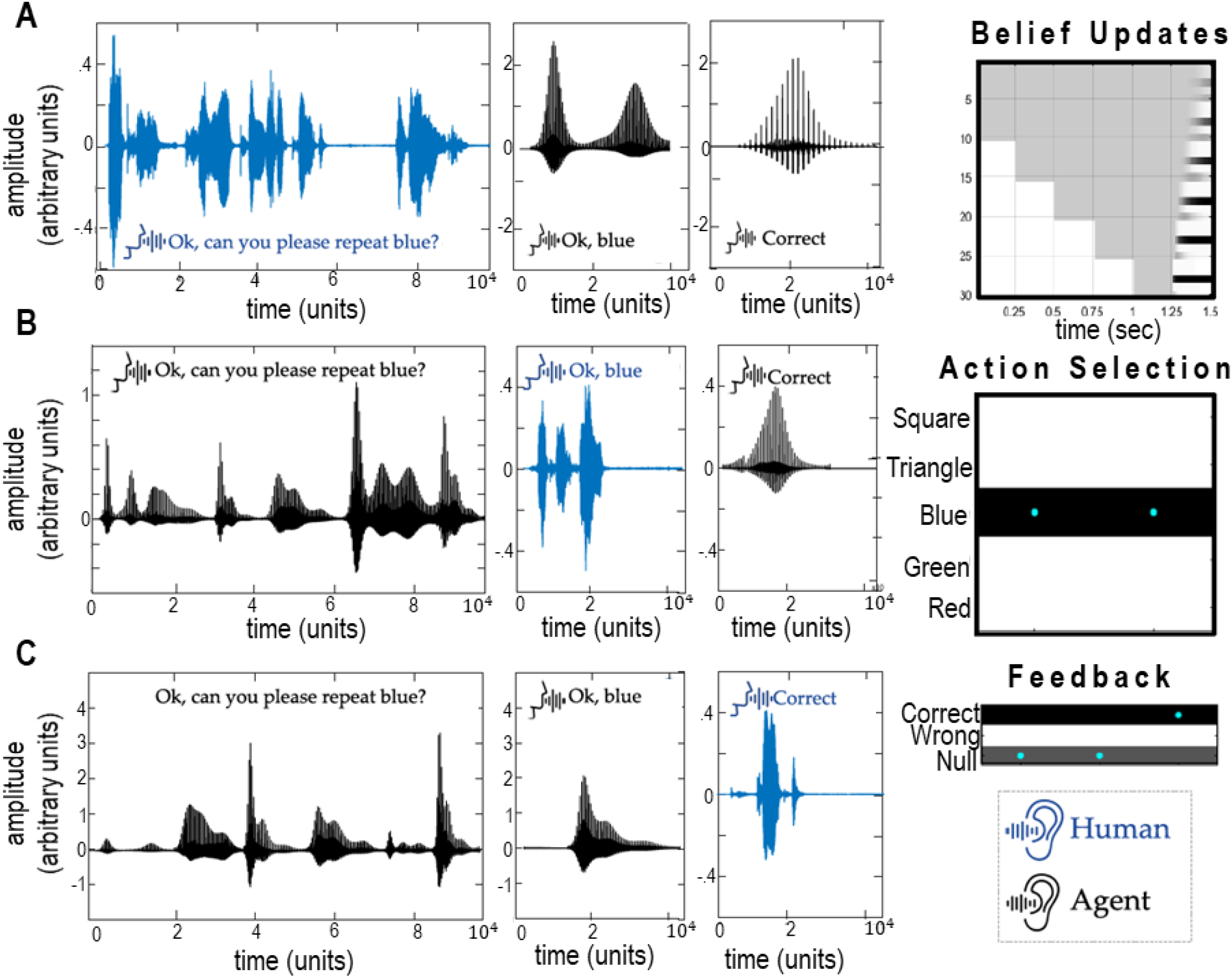
Human-in-the-loop simulations. These graphics illustrate the simulation results from one trial for when the real subject asks the question (ID A), repeats the target word (ID B) and gives feedback (ID C). For each row, the first three panels present the auditory waveform produced during the word repetition exchange. The waveform encodes the pattern of sound amplitude (y-axis) in the time domain (x-axis). Here, blue denotes the human subject, and black the agent. Notice that the waveform differs across the simulations (e.g., compare the first panel in rows ID A and ID B). This demonstrates WORM’s ability to effectively recognise and synthesise speech. The final panel in row ID A presents the belief updates for the *Target word* factor during the Question, equivalent to Figure 5B. The final panel in row ID B presents the selected action at the higher discrete level of the generative model. The x-axis represents time (for the first two timesteps when particular actions are selected) and y-axis displays the five possible actions. As in Figure 5, white is an expected probability of zero, black of one, and the cyan dots denotes the sampled action at each time. The final panel in row ID C presents the inferred feedback: the x-axis denotes time, and the y-axis shows the three *Feedback* options (Correct, Wrong, or Null). Again, white is an expected probability of zero, black of one, and cyan dots denote posterior beliefs.

## Discussion

In this paper, we have introduced a mixed generative model of word repetition called WORM. WORM has a hierarchical architecture with discrete higher levels that encode linguistic and lexical contexts mapping to a continuous auditory outcome (i.e., an auditory time series). This hierarchical organisation allows for a slowly evolving discrete level that contextualises turn-taking during the linguistic exchange, a faster discrete level that orders sequences of words into phrases, and fast continuous outcomes at the timescale of individual words. In WORM, we combined two distinct types of models. The higher levels were instantiated as partially observable Markov Decision processes (Figure 1) of the type previously used to simulate word repetition in (Sajid et al., 2020b; Sajid et al., 2020a; Sajid et al., 2021b). Conversely, a continuous model was used at the lower (faster) level: specifically, a generative model for synthesising and recognising speech (Friston et al., 2020b). By combining these formulations, the same model was used to: (i) recognise the linguistic context and target word from an acoustic signal, and (ii) generate the necessary acoustic stream, to repeat the heard word.

The belief update scheme, corresponding to variational inference, allowed for posterior evaluation of particular continuous signals—which we showed was possible based on the outputs from another WORM agent or a real human subject. WORM’s ability to appropriately communicate, and update beliefs, when interacting with human subjects speaks to its potential application in computational neuropsychology, specifically in identifying language disorders. Indeed, word repetition is a canonical paradigm in the neuropsychology of language (Swinburn et al., 2004). Future work could fit WORM to behavioural data collected from aphasic subjects (e.g., (Price et al., 2010; Seghier et al., 2016)), which could stratify different groups of subjects and provide predictions about corresponding changes in effective connectivity during the repetition of a heard word (Schwartenbeck and Friston, 2016). It might also be useful for simulating in-silico brain dysfunction to evaluate particular speech impediments.

Because we now have at hand a synthetic subject one can, in principle, examine the effects of synthetic lesions in different parts of the generative model. For example, one could reproduce the behaviours and responses characteristic of peripheral hearing loss by compromising the continuous dynamics involved in active listening; e.g., by blurring word boundaries, adding acoustic noise, or precluding the estimation of that noise. In a similar way, one could ask what would happen if the intrinsic connections that underwrite state transitions were compromised; for example, one could simulate the consequences of a failure to maintain the target word in working memory by appropriate lesions to the intrinsic connections (i.e., B-matrix) at the first discrete level of the model. Finally, one could simulate a failure to understand the structure of the paradigm, in terms of transitions at the higher discrete level that enable turn taking.

In a similar vein, interesting questions arise in relation to impairing the likelihood mappings (A) between different levels, and whether there will be distinct behavioural characteristics when (partially) disconnecting the implicit connections. In short, one can cast neuropsychology as pernicious disconnection syndromes and map out the distinct functional deficits that accompany specific disconnections or mappings that constitute the generative model.

The limitations of this work are self-evident; we used an amortised continuous generative model with learned empirical priors. Therefore, the model does not have capacity to make inferences over the word boundaries at the continuous level. Future work could combine the continuous and discrete models to allow for enactive processing of word boundaries. In other words, Bayesian beliefs about the current sequence of words at the discrete level would furnish empirical priors over alternative word segmentations or boundaries; thereby augmenting the robustness of word recognition to background noise or other distortions. Next, while we provide face validity, it is insufficient, at this stage, to demonstrate the applicability of this framework to large lexica in clinical applications. The scaling may require a lower-level generative model with amortised nonlinear mappings that are context insensitive—of the kind afforded by deep learning models (Yu and Deng, 2010; Tjandra et al., 2017). In future work, we will scale this model to include larger lexicon that allows for more elaborate exchanges beyond repeating heard words. For this, we can employ deep learning methods to amortise the mapping between the continuous auditory stream and discrete lexical, prosody and speaker states. In such instances, it will be important to determine whether recognition performance changes as a consequence of a larger lexicon.

Furthermore, coarticulation of words remains an issue in our model construction; this may be solved by incorporating priors forbidding the generation of multiple simultaneous words. Moreover, when considering how words are generated there is wide variability in the articulation of the same word across people (Hillenbrand et al., 1995; Remez, 2010) and—even when spoken by the same speaker—articulation depends on prosody (Bnziger and Scherer, 2005). From the perspective of recognition, two signals that are acoustically identical can be perceived as different words by human listeners, depending on their context; e.g., the preceding phonemes (Mann, 1980) or preceding spectral content (Holt et al., 2000). Future work could incorporate this kind of context sensitivity, in the lower level of the model, to allow for a more expressive form of active listening.

## Conclusion

In summary, this paper introduces a deep temporal model of word repetition (WORM) that generates a continuous acoustic signal in a linguistic exchange. The generated acoustic stream is sensitive to the turn-taking context and the target word to be repeated. The same model can also be used (in combination with a belief updating scheme) to infer a target word from a continuous acoustic signal. Incorporating turn-taking means that an agent who adopts this model knows when it is their turn to speak or listen. Through simulations (i.e., *in silico* ‘experiments’), we demonstrate that an agent equipped with this generative model can interact with another agent, with the same generative model, or with a real human subject— either taking on the role of the ‘experimenter’ or the ‘subject’ in a word repetition task. In future work, we plan to use WORM to simulate more complex exchanges and to make inferences about particular populations of (e.g., aphasic) subjects, paving the way towards further applications in computational neuropsychology.

## Software note

Although the generative model changes from application to application, the belief updates—and simulated neuronal responses—described in this paper are generic and can be implemented using standard routines (here spm_MDP_VB_X.m). These routines are available as Matlab code in the SPM academic software: http://www.fil.ion.ucl.ac.uk/spm/. The simulations in this paper can be reproduced (and customised) via the code and training data available at http://www.github.com/ucbtns/worm.

## Authors Contributions

All authors made substantial contributions to the conception, design and writing of the article; and approved publication of the final version.

